# Rapid development of cloud-native intelligent data pipelines for scientific data streams using the HASTE Toolkit

**DOI:** 10.1101/2020.09.13.274779

**Authors:** Ben Blamey, Salman Toor, Martin Dahlö, Håkan Wieslander, Philip J Harrison, Ida-Maria Sintorn, Alan Sabirsh, Carolina Wählby, Ola Spjuth, Andreas Hellander

## Abstract

This paper introduces the *HASTE Toolkit*, a cloud-native software toolkit capable of partitioning data streams in order to prioritize usage of limited resources. This in turn enables more efficient data-intensive experiments. We propose a model that introduces automated, autonomous decision making in data pipelines, such that a stream of data can be partitioned into a tiered or ordered *data hierarchy*. Importantly, the partitioning is online and based on data content rather than *a priori* metadata. At the core of the model are *interestingness functions* and *policies*. Interestingness functions assign a quantitative measure of interestingness to a single data object in the stream, an interestingness score. Based on this score, a policy guides decisions on how to prioritize computational resource usage for a given object. The HASTE Toolkit is a collection of tools to adapt data stream processing to this pipeline model. The result is smart data pipelines capable of effective or even optimal use of e.g. storage, compute and network bandwidth, to support experiments involving rapid processing of scientific data characterized by large individual data object sizes. We demonstrate the proposed model and our toolkit through two microscopy imaging case studies, each with their own interestingness functions, policies, and data hierarchies. The first deals with a high content screening experiment, where images are analyzed in an on-premise container cloud with the goal of prioritizing the images for storage and subsequent computation. The second considers edge processing of images for upload into the public cloud for a real-time control loop for a transmission electron microscope.

**Key Points:**

- We propose a pipeline model for building intelligent pipelines for streams, accounting for actual information content in data rather than *a priori* metadata, and present the HASTE Toolkit, a cloud-native software toolkit for supporting rapid development according to the proposed model.
- We demonstrate how the HASTE Toolkit enables intelligent resource optimization in two image analysis case studies based on a) high-content imaging and b) transmission electron microscopy.
- We highlight the challenges of storage, processing and transfer in streamed high volume, high velocity scientific data for both cloud and cloud-edge use cases.

## Introduction

Large datasets are both computationally and financially expensive to process, transport and store. Such datasets are ubiquitous throughout the life sciences, including imaging, where different types of microscopy are used to e.g. observe and quantify effects of drugs on cell morphology. Modern imaging techniques can generate image streams at rates of up to 1TB/hour [1]. Clearly, the processing, storing and communication of these images can be slow, resource-intensive and expensive, effectively becoming a bottleneck to scale experiments in support of data-driven life science. Another prominent example is human genome sequencing, where the global storage requirements is predicted to be between 2 and 40 exabytes (1 exabyte = 10^18^ bytes) by 2025, and with modern techniques generating data at the order of ~60 GB/h [2]. Similarly, in large-scale modeling, a single computational experiment in a systems biology context can generate terabytes of data [3].

This work is motivated by some of the most critical aspects of scalable scientific discovery for spatial and temporal image data. There are two primary concerns: (1) not all data is equally valuable. With datasets outgrowing resources, data storage should be prioritized for data that is most relevant (or interesting) for the study at hand and poor quality, or uninteresting, data (e.g. out-of-focus images) should be discarded or archived; (2) when resources are limited, or if decisions are required in real-time, we have to be smart about how the data is (pre)processed and which subsets of the data are stored for more detailed (and potentially computer intensive) analysis - prioritizing more interesting subsets of the data.

The general challenges of management and availability of large datasets are often popularized and summarized trough the so-called *Vs of big data*. Initially, the focus was on the three Vs: velocity, volume and variety, but this list has since grown with the increasing number of new use-cases to also include *Vs* such as veracity, variability, virtualization and value [4]. During the last decade, a number of frameworks have been designed to address these challenges, offering reliable, efficient and secure large-scale data management solutions. However, according to a white paper published by IDC [5], only 30% of the generated data is in the form that it can be efficiently analyzed. This highlights the current gap between large-scale data management and efficient data analysis. To close this gap, it is essential to design and develop intelligent data management frameworks that can help organize the available datasets for efficient analyses. In this work, we address this challenge by proposing a model that helps a data pipeline developer make online decisions about individual data objects’^1^ priority based in actual information content, or interestingness, rather than traditional metadata.

A range of existing work in life science applications has discussed the challenges of transporting, storing and analyzing data, often advocating a streamed approach. In [6] the authors explicitly discuss the constraints of cloud upload bandwidth, and its effect on overall throughput for mass-spectrometry based metabolomics. In their application, uploading large datasets from the instrument to the cloud represents a bottleneck and they advocate a stream-based approach with online analysis where data is processed when it arrives, rather than waiting for upload of the complete dataset. Hillman et al. [7] developed a stream based pipeline with Apache Flink and Kafka for processing of proteomics data from liquid chromatography-mass spectrometry (LC/MS) and note the advantages of a realtime approach to analysis: “a scientist could see what is happening in real-time and possibly stop a problematic experiment to save time”. Zhang et al. [8] developed a client/server application for interactive visualization of MS spectra, adopting a stream-based approach to achieve better user interactivity. In genomics, [9] presented the htsget protocol to enable clients to download genomic data in a more fine-grained fashion, and allow for processing chunks as they come from the sequencer. In [10], the authors note that a single electron microscope can produce 1 TB of images per day, requiring a minimum of 1000 CPU hours for analysis. Adapting their Scipion software [11] (intended for Cryo EM image analysis) for use in the cloud, they discuss the challenges of data transfer to/from the cloud, comparing transfer rates for different providers. [12] proposes excluding outliers in streaming data, using an ‘Outlier Detection and Removal’ (ODR) algorithm which they evaluate on five bioinformatics datasets.

Rather than handling one particular type of data or dealing with a specific data pipeline, the aim of the present work is to distill effective architectural patterns into a pipeline model to allow for repeatable implementations of smart systems capable of online resource prioritization in scenarios involving large-scale data production, such as from a scientific instrument. Computers in the lab connected directly to such an instrument, used together with cloud resources, are an example of edge computing [13]. Under that paradigm, computational resources outside the cloud (such as mobile devices, and more conventional compute nodes) are used in conjunction with cloud computing resources to deliver benefits to an application such as reduced cost, better performance, or improved user experience. General computer science challenges include security, deployment, software complexity, and resource management/workload allocation. In our context, the streams of large data objects generated by scientific instruments create particular challenges within the edge computing paradigm as the data often needs to be uploaded to the cloud for processing, storage, or wider distribution. Whilst limited compute resources at the edge are often insufficient for low-latency processing of these datasets, intelligent workload allocation can improve throughput (as discussed in Case Study 2).

In this paper we propose a pipeline model for partitioning and prioritizing stream datasets into *data hierarchies* (DHs) according to an *interestingness function* (IF) and accompanying *policy*, applied to objects in the stream, for more effective use of hardware (in edge and cloud contexts). We present this as a general approach to mitigating resource management challenges, with a focus on image data. Our model allows for autonomous decision making, while providing a clear model for domain experts to manage the resources in distributed systems - by encoding domain expertise via the IF. To that end, this paper introduces the HASTE^2^ Toolkit, intended for developing intelligent stream processing pipelines based on this model. Two case studies presented in this paper document how microscopy pipelines can be adapted to the HASTE pipeline model.

Whilst the core ideas of intelligent data pipelines in HASTE is generally applicable to many scenarios involving scientific datasets, we here focus on case-studies involving image streams, in particular from microscopy.

## Background: Stream Processing and Workflow Management

A fundamental component to the pipelines presented in this paper is a stream processing engine. Systems for stream processing are generally concerned with high frequency message influx, and those objects can be small in size, such as a few KB. Examples of such data objects include sensor readings from IoT devices (such as MQTT messages), those generated from telecoms, web and cloud applications, e-commerce and financial applications, or the aggregation and analysis log entries. Well-known examples of mature, enterprise-grade frameworks for cloud-based stream processing in these contexts include Apache Flink, Apache Spark Streaming, and Apache Log Flume. Resilience and fault tolerance are key features of these frameworks (often achieved with various forms of redundancy and replication). These frameworks are commonly used in conjunction with various queuing applications, e.g., Apache Kafka, and vendor-specific products such as AWS Kinesis - these also include basic processing functionality.

Whilst the maturity, support, documentation, features and performance (order of MHz message processing throughput) boasted by these frameworks is attractive for scientific computing application, streamed scientific data (and its processing) tends to have different characteristics: data objects used in scientific computing applications (such as microscopy images, and matrices from other scientific computing domains) can be larger in size, which can create performance issues when integrated with these enterprise frameworks described above [14]. For example, data object sizes in imaging applications could be a few MB.

To address this gap, we have previously developed and reported on a stream processing framework focusing on scientific computing applications, HarmonicIO [15]. HarmonicIO sacrifices some of these features, and is intended for lower-frequency applications (towards kHz, not MHz), and was able to achieve better streaming performance under some conditions in one study for larger message sizes [14]. The HASTE Toolkit has been developed with images as the primary usecase and for this reason HarmonicIO is the default supported streaming framework in the toolkit. However, we stress here that in principle any streaming framework can be used.

Furthermore, under the emerging edge computing paradigm, there are some stream processing frameworks available, often focusing on traditional IoT use-cases. Being in their infancy, effective automated scheduling and operator placement in hybrid edge/cloud deployment scenarios remains an open research challenge for this context. Within this area, there is significant research effort concerning real-time video analysis, where images collected at the edge (from cameras) are streamed to the cloud for analysis - some degree of lossy compression is typically used in such applications.

By contrast, *workflow* frameworks are broad class of software frameworks intended to facilitate the development of data-processing pipelines. There are a large number of such frameworks (more than 100 are listed in [16]). In such frameworks, one generally defines processing operations (often as the invocation of external processes), which are triggered by events such as the creation of a new file on disk, or a commit being pushed to a Git repository. Such frameworks generally handle large numbers of files, of arbitrary size, and often include some degree of fault tolerance. But in contrast to stream processing frameworks, they may lack functionality specific for streams, such as window operations, more complex scheduling and placing of operators, and are generally intended for higher latency and/or lower ingress rates (than the 100kHz+ range of the stream processing frameworks described above), and are often file-system centric, with objects being written back to disk between each processing step.

The HASTE toolkit attempts to fill a gap between these two classes of software *(stream processing frameworks*, and *workflow management systems*): applications where latency and high data object throughput are important (and use of a filesystem as a queuing platform are perhaps unsuitable for that reason), but not as high as some enterprise stream processing applications; whilst being flexible enough to accommodate a broad range of integration approaches, processing steps with external tools, and the large message sizes characteristic of scientific computing applications.

The priority-driven approach of the HASTE pipeline model reconciles the resource requirements of life science pipelines (characterised by streams relatively of large message, with expensive per-message processing steps), with the requirements for low-latency and high throughput, allowing for real time human supervision, inspection, interactive analysis - as well as real-time control of laboratory equipment.

## HASTE Pipeline Model

The key ideas of the HASTE pipeline model are the use of *interestingness* functions and a *policy* to autonomously induce *data hierarchies*. These structures are then used to manage and optimize different objectives such as communication, processing and storage of the datasets. The HASTE Toolkit enables rapid constructions of smart pipelines following this model. Central to the approach is that decisions are made based on actual data content rather than on *a priori* metadata associated with the data objects. The following subsections introduces the components of the pipeline model.

### Overview

Figure 1 illustrates the proposed HASTE model and logical architecture. One or more streaming data sources generate streams of data objects. The stream then undergoes feature extraction (relevant to the context) - this data extraction can be performed in parallel, as an idempotent function of a single object. The intention is that computationally cheap initial feature extraction can be used to prioritize subsequent, more expensive, downstream processing.

An Interestingness Function (IF) computes an interestingness score for each object from these extracted features. This computation can be a simple procedure, e.g. to nominate one of the extracted features as the interestingness score associated with the data object. In more complex cases it can also be a machine learning model trained either before the experiment or online during the experiment that generates the stream. Finally, a policy is applied which determines where to store the object within a Data Hierarchy (DH), or send it for further downstream processing, based on the interestingness scores.

**Figure 1.**
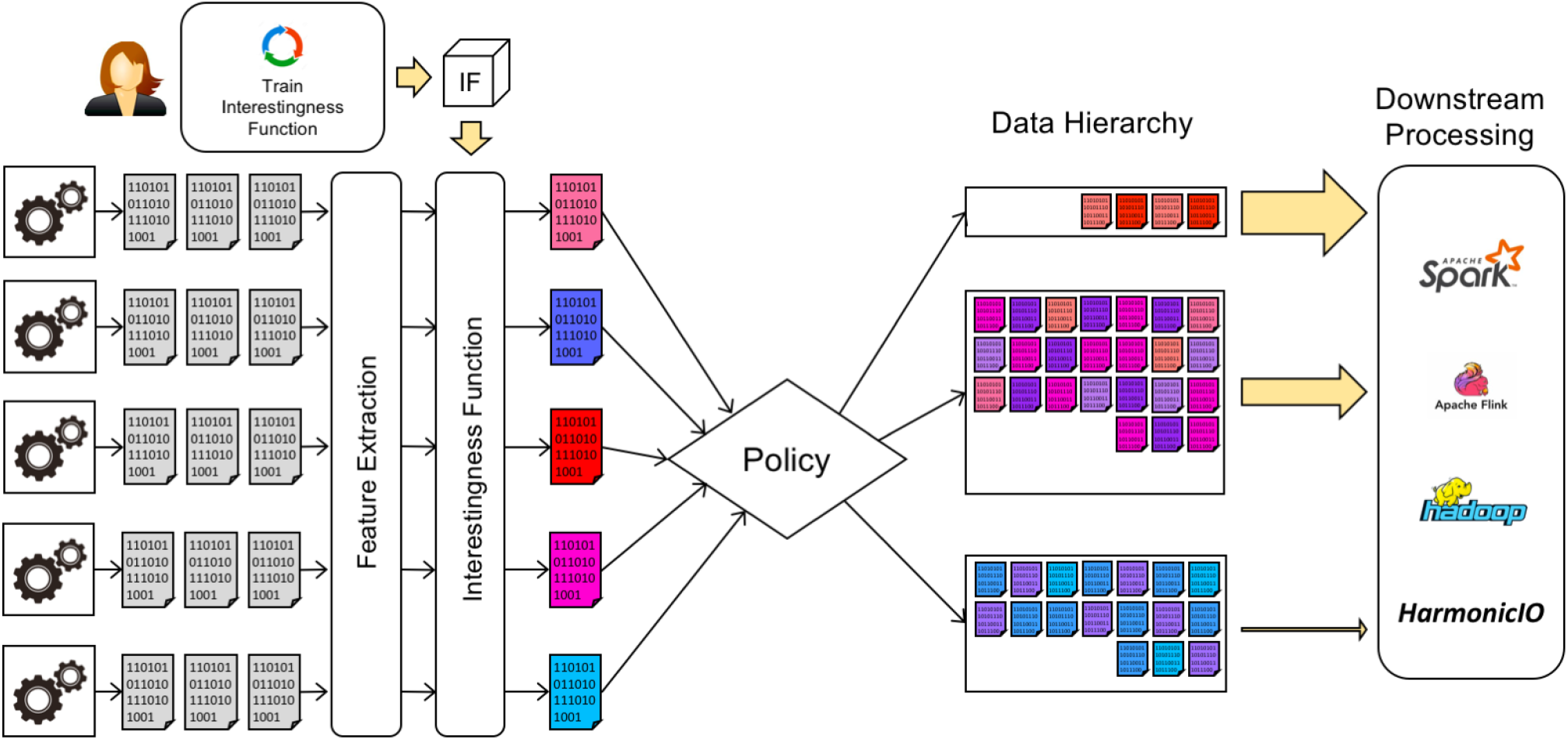
Logical Architecture for the HASTE pipeline model. A stream of data objects is generated by one or more streaming sources (such as a microscope). These objects undergo online, automated feature extraction, and an IF is applied with the extracted features as input. This associates an interestingness score with each object in the stream. A user-defined policy is then used to organize the data objects into a data hierarchy to be used for optimizing subsequent communication, storage and downstream processing.

### Interestingess Functions

The IF is a user provided function, to be applied to the extracted features from the data objects in the stream. The purpose of the IF is to associate an *interestingness score* with each object. Examples of IFs in image analysis contexts could be features relating to image quality, detected phenomena in images, etc.

The computed IF score is used for determining priority for subsequent processing, communication, and/or storage of that object. In this sense, IFs have some similarities to the concept of document (data object) ‘hotness’ in tiered storage contexts, where a more recently accessed ‘hot’ document would be stored in a high-performance tier. Whilst much of that line of work uses only file-system information, other work takes some consideration of the application itself, for example [17] model access as a Zipf distribution, for a review see [3].

Our present work generalizes the concept of ‘hotness’, in a number of ways: (1) our IFs always take consideration of semantics at the level of the scientific application – in the case of microscopy imaging this could be image focus, or quality features - perhaps combined with business logic for particular color channels, etc. - rather than file system semantics (such as file access history). This approach allows an immediate, online decision about the object’s interestingness - rather then inferring it from subsequent access patterns. (2) Tiered storage is just one potential application of HASTE: we use IFs to prioritize data objects for storage, compute, and communication (3) with HASTE, the intention is that users’ configure IFs themselves, together with the associated policy. Currently, the output of the IF is scalar valued. This is intended to assure smooth integration in cases where the IF is a machine learnt model, outputting a probability, rank, or some other statistical measure.

Further, we propose general software abstractions for these ideas, and demonstrate the potential benefits of online interestingness-based prioritization in two case studies: both in terms of the optimization of various resources (compute, communication, storage), but also from an experimental and scientific viewpoint - selecting the best data (or outliers) for inspection and further analysis.

### Policies for inducing Data Hierarchies

In applications utilizing tiered storage, more interesting data objects would be saved in higher performance, more expensive tiers - readily accessible for downstream processing (while less interesting objects could be cheaply archived) - explored in Case Study 1. Whereas, in an edge computing contexts, we may want to prioritize data objects for computation at the cloud edge, to make more effective use of that resource - explored in Case Study 2. In both cases we refer to these structures as *data hierarchies* (DHs). In a HASTE pipeline DHs are ‘induced’ within the source data by the IF and a policy. The *policy* takes the interestingness score as input and applies a set of rules to determine how an object is placed within the DH, for example, its allocation within a tiered storage system; or where it should be stored or processed downstream. Listing 1 shows how a user can define a policy, a simple dictionary mapping intervals of interestingness scores to the tiers (which are configured separately). In this paper we demonstrate two forms of policy: the interval model mentioned above (where the tier is determined directly from the interestingness score, Case Study 1) and a priority-based policy, where data objects are queued (according to their interestingness) for upload and processing (as in Case Study 2).

A benefit of the HASTE pipeline model is the clear role separation - all the domain-specific knowledge is effectively encapsulated within the IF whilst the choice of how to form DHs and optimize storage tier allocation is encapsulated entirely within the policy. This allows the scientific question of what constitutes an interesting data object, and the computing infrastructure, or indeed, budgetary, concerns of how to make best use of computing resources (including storage), to be separated and worked on by team members with different expertise. Importantly, this de-coupling allows the possibility for IFs to be reused among scientists, and between contexts where the data may be similar, but the dataset size, and available computing infrastructure, may be different.

## The HASTE Toolkit

The HASTE Toolkit implements the core functionality needed for rapidly constructing smart pipelines based on the proposed model.

### HASTE Storage Client

The HASTE Storage Client (HSC) serves as the main entrypoint for the user. It is configured with the IF, the policy, and the configuration associated with the tiers, and processes each data object arriving in the stream. It can be installed as a standalone Python module (see: https://github.com/HASTE-project/HasteStorageClient, version 0.13 was used for this study.). It allows a DH to be realized within HASTE as tiered storage. It is a library with the core prioritization functionality: it invokes the IF on incoming data objects, and applies the policy to form the data hierarchy. The extracted features are used to compute interestingness scores, along with other metadata and logging info, are saved in a database by the HSC. It is intended to be adopted within the Python-based stream processing framework of choice, an example can be found at: https://github.com/HASTE-project/HasteStorageClient/blob/master/example.py. An existing pipeline can be adapted to use HASTE according to the following steps:

- Install the HSC from PyPI `pip install haste-storage-client`, or from source.
- Configure one or more storage tiers (on a HASTE-compatible storage platform)^3^.
- Define an IF for the context - it can use spatial, temporal or other metadata associated with the data object.
- Run feature extraction on the object prior to invoking the HSC.
- Deploy a MongoDB instance. The scripts https://github.com/HASTE-project/k8s-deployments/ can be adapted for this purpose.

### Other key components of the HASTE Toolkit

This section lists other various components in the HASTE toolkit, and describes how they relate to the key ideas of IFs, data hierarchies (DHs) and policies.

#### The HASTE Agent

A command-line application (developed for the microscopy use case in Case Study 2), which uploads new documents on disk to the cloud, whilst performing intelligently prioritized pre-processing of objects waiting to be uploaded, so as to minimize the overall upload duration. (see: https://github.com/HASTE-project/haste-agent (v0.1 was used for this study)). The functionality of this tool is discussed in detail in Case Study 2.

#### The HASTE Gateway

Cloud gateway service, which receives data objects in the cloud, and forwards them for further processing. Deployed as a Docker container. (see: https://github.com/HASTE-project/haste-gateway, v0.1 was used in this study.).

#### The HASTE Report Generator

An auxiliary command line tool for exporting data from the **Extracted Features Database**. (see: https://github.com/HASTE-project/haste-report-generator).

#### The Extracted Features Database

MongoDB is used by the **HASTE Storage Client** to hold a variety of the metadata: extracted features, interestingness scores, and tier/DH allocation.

#### Tiered Storage

Tiered storage is one way that a data hierarchy (DH) can be realized. The HSC allows existing storage to be organized into a tiered storage system, where tiers using various drivers built into the HSC can be configured. In Case Study 2 the tiers are filesystem directories, into which image files are binned according to the user-defined policy. The idea is that in other deployments, less expensive disks/cloud storage could be used for less interesting data. Note that the policy can also send data deemed unusable (e.g. quality below a certain threshold) directly to trash. Tiered storage drivers are managed by the HASTE storage client.

Our GitHub project page (https://github.com/HASTE-project) showcases other components relating to various example pipelines developed within the HASTE project, including IFs developed for specific use cases as well as scripts for automated deployment.

## Experiments and Results

In this section we illustrate the utility of the toolkit in two real-world case studies chosen to demonstrate how the HASTE pipeline model can be realized in practice to optimize resource usage in two very different infrastructure and deployment scenarios. Table 1 summarizes the objectives of the case studies.

**Table 1.** Overview of the two case studies used in this paper.

|  | Case Study 1 – Cell Profiling | Case Study 2 – Real Time Processing with a TEM |
| --- | --- | --- |
| Application | High-Content Imaging | Real Time Control of Microscopy |
| Prioritization of... | Storage | Communication & Compute |
| Goal | Tiered Storage | Reduce end-to-end latency for cloud upload |
| Deployment Setting | On-premises Cloud, Kubernetes | Cloud Edge & Public Cloud (SNIC) |
| Interestingness Function (IF) | CellProfiler Pipeline – Image Quality | Estimation of Size Reduction (Sampling, Splines) |
| Policy | Fixed Interestingness Thresholds | Dynamic Interestingness Rank |

Case Study 1 concerns data management for a high-content screening experiment in a scientific laboratory at Uppsala University, Sweden. A small on-premises compute cluster running Kubernetes [19] provides the necessary infrastructure to handle the immediate data flow from the experiment, but both storage capacity and manual downstream analysis is a concern. We use the HASTE toolkit to build a pipeline that captures the input data as an image stream and bins images into tiers according to image quality. The overall goal is to organize the images into tiers for subsequent processing, to both ensure that the best images are allocated to high performance storage for high performance analysis, and to help the scientist prioritize manual work to appropriate subsets of data.

Case Study 2 concerns processing an image stream from a transmission electron microscope (TEM). During streaming, there is an opportunity to pre-process images using a desktop PC co-located with the microscope, before being uploaded for stream processing in the cloud. This is an example of an edge computing [13]) scenario, where very limited but low-latency local infrastructure is leveraged together with a large cloud infrastructure. The ultimate goal is real-time control of the microscope (see Figure 6), and consequently end-to-end latency is a key concern. This latency is constrained by image upload time. Here we develop a pipeline using the HASTE tools with an IF that predicts the effectiveness of pre-processing individual images at the edge prior to cloud upload.

### Case Study 1 – Smart data management for high-content imaging experiments

This case study focuses on adoption of the HASTE toolkit in a high-content microscopy setting – the input is a stream of images arriving from an automated microscope. This deployment uses an on-premises compute cluster running Kubernetes with a local NAS. While we want online analysis, we consider this a ‘high latency’ application - images can remain unprocessed for some seconds or minutes until compute resources are available. This is a contrast to Case Study 2, where low-latency processing is a goal.

Image quality is an issue in microscopy: images that have debris, are out of focus, or unusable for some other reason relating to the experimental setup. Such images can disrupt subsequent automated analysis and are distracting for human inspection. Furthermore, their storage, computation and transportation have avoidable performance and financial costs.

For this case study, the HASTE toolkit is used to prioritize storage. The developed IF is a CellProfiler pipeline performing *out of focus* prediction using the `imagequality` plugin [20]). The *Policy* is a fixed threshold used to bin images into a DH according to image quality. See Table 1 for an overview of the case studies.

Figure 2 illustrates the key components of the architecture:

- Client – monitors the source directory for new image files, adding the name of each file to the queue. (see: https://github.com/HASTE-project/cellprofiler-pipeline/tree/master/client, v3 was used for this study).
- Queue – a RabbitMQ queue to store filenames (and associated metadata). Version 3.7.15 was used for this study.
- Worker – waits for a filename message (on the queue), runs a CellProfiler pipeline on it, computes an interestingness score from the CellProfiler features (according to a user-defined function). (see: https://github.com/HASTE-project/cellprofiler-pipeline/tree/master/worker, v3 was used for this study)

**Figure 2.**
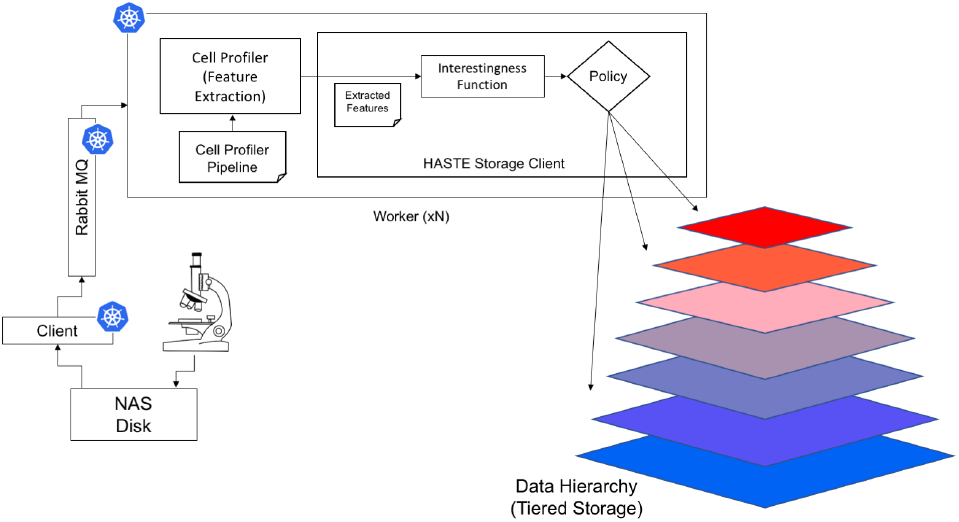
Architecture for Case Study 1. In this case study, the DH is realized as storage tiers. Images streamed from the microscope are saved to disk (Network Attached Storage). This disk is polled by the ‘client’, which pushes a message about the new file to RabbitMQ. Workers pop these messages from the queue, analyze the image, and move it to the storage tiers configured in the data hierarchy, using the HASTE Storage Client, configured with an appropriate IF and Policy. Icons indicate the components running as Kubernetes *pods*.

The deployment scripts for Kubernetes & Helm used to deploy these services for this study are available at: https://github.com/HASTE-project/k8s-deployments, v1.1 was used.

The image; together with its interestingness score and metadata are passed to the **HASTE Storage Client** – which allocates the images to **Tiered Storage/DH**, and saves metadata in the the **Extracted Feature Database**. Each image is processed independently, which simplifies scaling.

The HASTE toolkit simplifies the development, deployment and configuration of this pipeline – in particular, the interaction between the filesystems used in the input image stream and archive of the processed images. When using our image processing pipeline, user effort is focused on (a) defining a suitable IF and (b) defining a policy which determines how the output of that function relates to DH allocation (storage tiers). Both of these are declared within the Kubernetes deployment script. When developing the pipeline itself, one is able to provide the interestingness score (the output of the IF), and the policy as arguments to the HASTE tools, and delegate responsibility to applying the policy (with respect to the storage tiers), recording all associated metadata to the **Extracted Feature Database**.

The client, queue and workers are all deployed in Docker containers. Auto-scaling is configured for the workers: they are scaled up when processing images, and scaled back down again when idle. A message containing the image filenames (and other metadata) is queued, but the file content is read from the NAS for processing and tiering.

The code for the worker is an extension of *Distributed-CellProfiler* (released as part of CellProfiler v3.0) [21]^4^, which it to run within AWS^5^. The key benefit of our containerized system is that because it runs in Docker, and is not dependent on AWS services, it can be used for local deployments in laboratory settings, so that images do not need to be uploaded to the public cloud for processing. Alternatively, our system can be used with any cloud computing provider able to host Docker containers. We use the open-source message broker RabbitMQ in place of Amazon SQS (simple queue service). Our Kubernetes deployment scripts handle the necessary configuration, and a benefit of RabbitMQ is that it has a built in management web GUI. A helper script is provided to configure the credentials for the management GUI.

#### Evaluation

For validation of this case study we simulated analysis and tiering using a high content screening dataset previously collected in the lab, consisting of 2699 images of cortical neuronal cells, imaged with an ImageXpress XLS, the dataset is available at [22]. In doing so, we demonstrate that our system is able to handle a large number of images. To simulate the microscope, the images were copied into the source directory, triggering messages from the client, which were read by workers to analyze the images (with CellProfiler) to extract the relevant features from the results, apply the IF, and allocate them to the tiers according to the policy. Running in our laboratory Kubernetes environment, 17 workers were able to process images simultaneously.

We use the PLLS (Power Log Log Slope) feature as the basis of our interestingness score, as it has been shown to be a robust measure of image focus [23]. In this case study, we use the logistic function *f* as an IF, applying it to the PLLS feature *x*, to compute the interestingness score. The logistic function has output in the range (0,1):

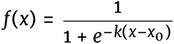

The PLLS values will depend on a number of factors (such as magnification, number of cells, stainings, exposure times, etc.). The parameters of this IF can be chosen to fit the modality, based on a sample of pre-images for calibration. In this case, we chose (*k* = 4.5, *x*_0_ = −1.4). The policy is defined to map the interestingness score in the intervals (*i*/4, (*i* + 1)/4) for *i* ∈ (0,1,2,3) to the respective storage tiers. Figure 3 shows histograms of the PLLS feature and Interestingness Score.

**Figure 3.**
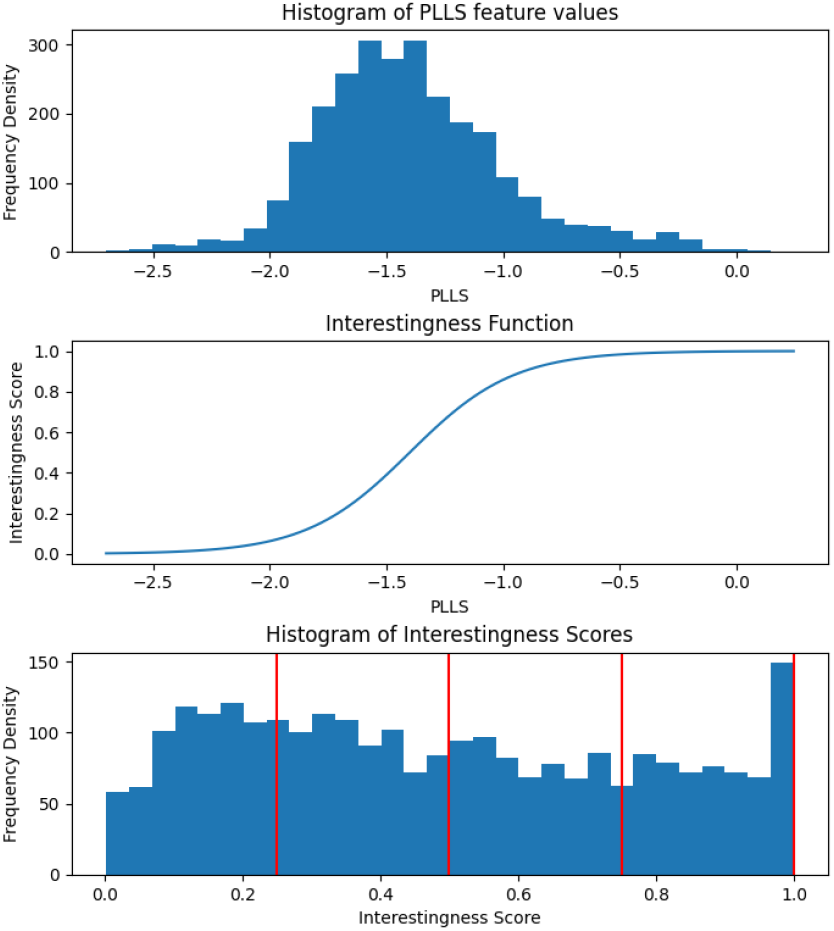
Histograms of the PLLS feature scores (top), and when converted to an Interestingness Score (bottom), by application of the Logistic Function (the IF for Case Study 1, middle). The vertical lines on the bottom plot indicate tier boundaries configured in the policy. c.f. example images in Figure 4

For this evaluation, these tiers were simply directories on disk. Any storage system compatible with the HASTE Storage Client could be used, the key idea is that different storage platforms (with different performance and cost) can be used for the different tiers. In this case, we simply use the tiers as a convenient way to partition the dataset for further analysis and inspection. Figure 4 shows examples of the images according to tiers, and Table 2 shows the results.

**Figure 4.**
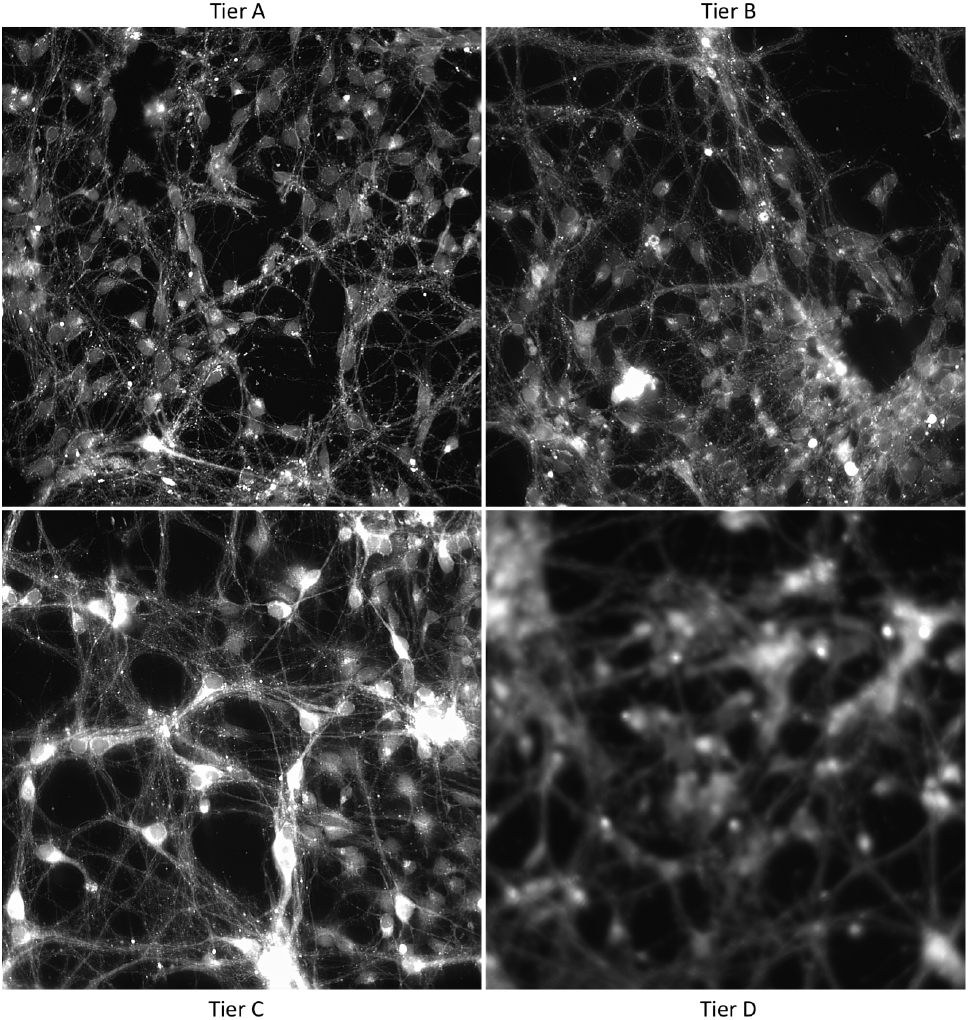
Example images from the high content screening dataset (Case Study 1), according to automatically assigned tier. Tier A is the most in-focus, with the highest PLLS feature values and interestingness scores.

**Table 2.**
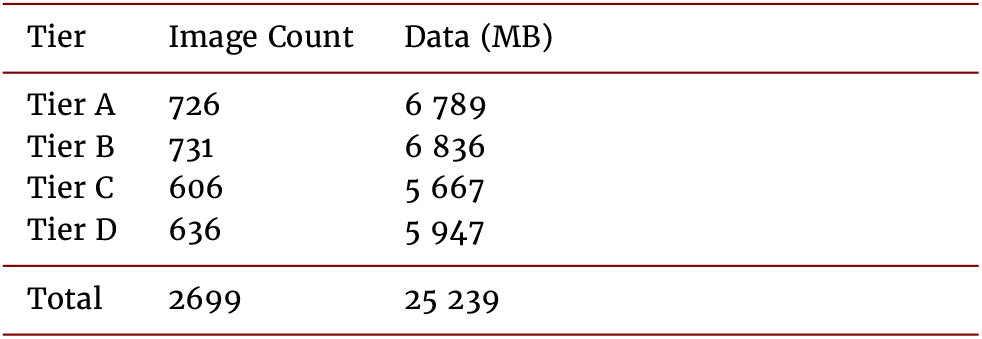
Image allocation for Case Study 1.

| Tier | Image Count | Data (MB) |
| --- | --- | --- |
| Tier A | 726 | 6 789 |
| Tier B | 731 | 6 836 |
| Tier C | 606 | 5 667 |
| Tier D | 636 | 5 947 |
| Total | 2699 | 25 239 |

~~~
interestingness_function(features):
     plls = features[’PLLS’]
     int_score = 1/(1 + exp(−(4.5) * (plls - (−1.4))))
     return int_score
See: https://github.com/HASTE-project/cellprofiler-pipeline/blob/master/worker/haste/pipeline/worker/LogisticInterestingnessModel.py
storage_policy:
[ [0., 0.25, tierD],
  [0.25, 0.50, tierC],
  [0.50, 0.75, tierB],
  [0.75, 1.00, tierA] ]
See: https://github.com/HASTE-project/k8s-deployments/blob/master/pipeline_worker.yaml
~~~

Listing 1: Pseudocode for Image Tier Placement (Case Study 1). The IF is the logistic function, applied to the previously extracted PLLS feature. The policy shows thresholds for the different tiers.

### Case Study 2 – Prioritizing analysis of TEM images at the Cloud Edge

This case study is concerned with the prioritized processing of a stream of images from a microscope (according to an IF), applied to a hybrid edge/cloud stream processing deployment context. In this example, we show how the HASTE tools can facilitate a better use of constrained upload bandwidth and edge compute resources. The image stream comes from MiniTEM^™^ - a 25keV transmission electron microscope [24] (Vironova, Sweden), connected to a desktop PC from which the microscope is operated and the image stream received, via proprietary driver software. The stream processing application preprocesses the TEM images locally (i.e. at the cloud edge), to reduce their image size, with the effect of reducing their upload time to the cloud, and hence the end-to-end processing latency.

The purpose of the pipeline is to automate a typical workflow for TEM analysis, which proceeds as follows: a sample is loaded into the microscope (in this case a tissue sample), the operator performs an ‘initial sweep’ over the sample at low magnification, to locate target (i.e. interesting) regions of the sample. In the conventional workflow, the search for ‘target’ areas of the sample is done by human inspection. The operator then images identified target areas of the sample at higher magnification for subsequent visual/digital analysis.

Automating this process entails the detection of target regions of the sample using an automated image processing pipeline, based on a set of images from the initial sweep. Such a pipeline would output machine-readable instructions to direct the microscope to perform the high magnification imaging, reducing the need for human supervision of sample imaging. The image processing pipeline used to detect target regions can be costly and slow and could hence preferably be performed in the cloud. Performing image processing in the cloud has several advantages: it allows short-term rental of computing resources without incurring the costs associated with up-front hardware investment and on-premises management of hardware. Machines with GPUs for deep learning, as well as secure, backed-up storage of images in the cloud, are available according to a pay-per-use model. With our overall aim of supporting a real-time control loop, and given the expense of the equipment, sample throughput is important. Despite images being compressed as PNGs, upload bandwidth is a bottleneck. Note that PNG compression is lossless, so as not to interfere with subsequent image analysis. Consequently, we wish to upload all the images from the ‘initial sweep’ into the cloud as quickly as possible, and this is what is targeted here.

A pre-processing operator, would reduce the compressed image size to an extent depending on the image content. However, this operator itself has a computational cost but because of the temporary backlog of images waiting to be uploaded, there is an opportunity to pre-process some of the waiting images to reduce their size (see Figure 5). The available upload bandwidth with respect to the computational cost of the pre-processing operator, means that (in our experiment) there is insufficient time to pre-process all images prior to upload. In fact, to pre-process all of them would actually increase end-to-end latency, due to the computational cost of the preprocessing operation and limited file size reduction for some images (content dependent). The solution is to prioritize images for upload and pre-processing respectively, whilst both processes, as well as the enqueuing of new images from the microscope, are occurring concurrently.

**Figure 5.**
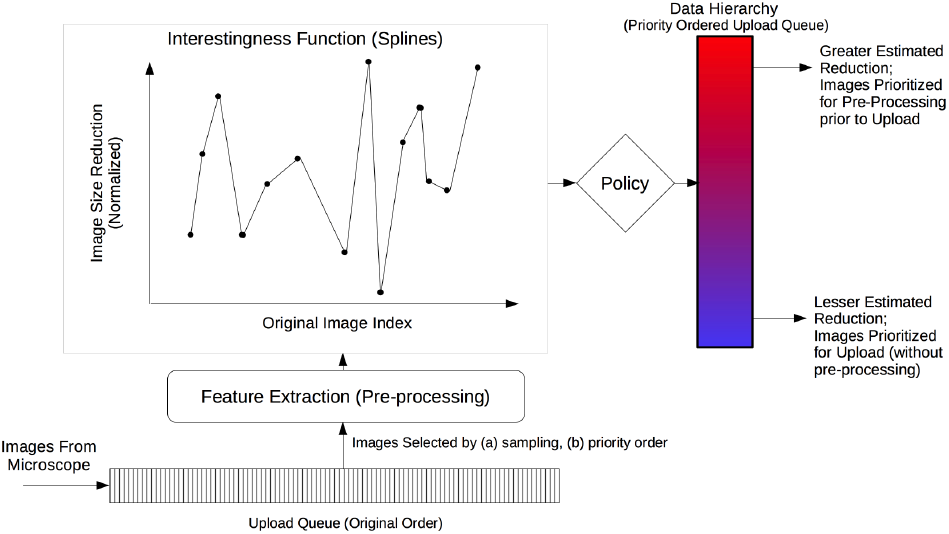
Architecture for Case Study 2, showing internal functionality of the Haste Desktop Agent at the cloud edge. Images streamed from the microscope are queued at the edge for uploading after (potential) pre-processing. The DH is realized as a priority queue. Images are prioritized in this queue depending on the IF which estimates the extent of their size reduction under this preprocessing operator: those with a greater estimated reduction are prioritized for processing (vice-versa for upload). This estimate is calculated by interpolating the reduction achieved in nearby images (see Figure 7). This estimated spline is the IF for this case study.

**Figure 6.**
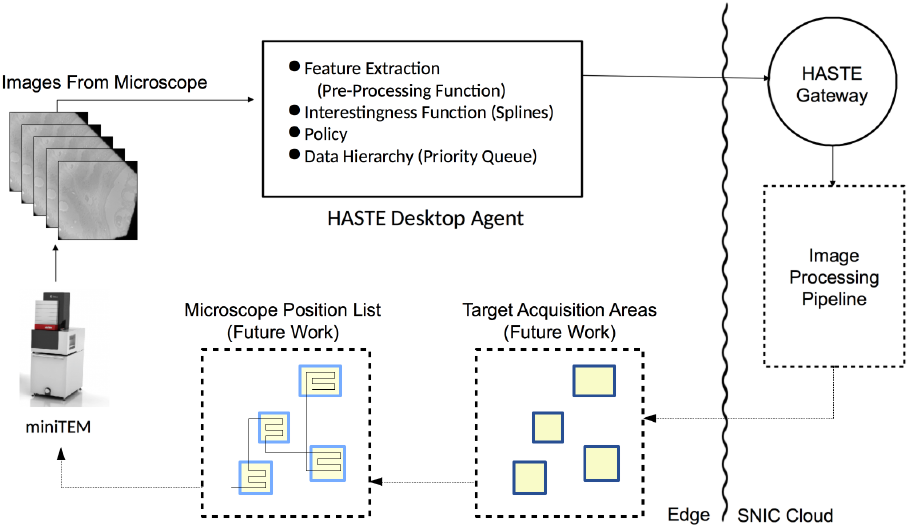
Architecture of the intended application: full control loop for the MiniTEM, with automatic imaging of target areas identified in initial scan. Control of microscope acquisition is future work. The internals of the HASTE Desktop Agent (where the HASTE model is applied) are shown in Figure 5.

#### Feature Extraction, the Interestingness Function, and Policy

Samples for TEM analysis are typically supported by a metal grid, which then obscures (blocks) regions of the sample in (in this case) a honeycomb pattern. The blocked regions appear black in the images. As the sample holder moves under the camera, the extent to which the sample is obscured is a piecewise smooth (but irregular) function of document index, dependent on the particular magnification level, and speed and direction of the sample holder movement. Images can be preprocessed to remove noise from blocked regions of the image, reducing the size of the image under PNG compression. The extent of file size reduction (under our pre-processing operator) is related to the extent to which the grid obscures the image.

Consequently, the predicted extent of file size reduction can be modelled with linear spline interpolation, based on the actual file size reduction of images sampled from the queue, described in more detail in [25]. The file size reduction corresponds to feature extraction in the HASTE pipeline model, and the spline estimate – the estimate of message size reduction – can be encapsulated as an IF, see Figure 1. The HASTE tools, specifically the **HASTE Agent** allow that IF to be used as a scheduling heuristic to prioritize upload and local (pre-)processing respectively (i.e. corresponding to the policy inducing the DH in HASTE).

Available compute resource at the cloud edge are prioritized on those images expected to yield the greatest reduction in file size (normalized by the compute cost, i.e. CPU time, incurred in doing so). Conversely, upload bandwidth is prioritized on (a) images that have been processed in this way, followed by (b) those images for which the extent of file size reduction is expected to be the least – under the aim of minimizing the overall upload time.

An important distinction between the this setting and that in Case Study 1 is that the IF and DH are dynamic in this case study.

The **HASTE Agent** manages the 3 processes occurring simultaneously: new images are arriving from the microscope, images are being pre-processed, and images are being uploaded.

#### Evaluation

When evaluated on a set of kidney tissue sample images [26] our edge-based processing approach yielded up to a 25% reduction in end-to-end stream processing latency (for this particular choice of processing operator and dataset) compared to a baseline approach without any prioritization, when compared to performing no stream processing at all [25]. This is a signifi-cant gain obtained with relative ease due to the HASTE Toolkit.

To verify the pre-processing operator, it was applied to all images after the live test was performed. Figure 7, shows how the image size reduction (y-axis – normalized with computational cost) can be modelled as a smooth function of the document index (x-axis). The colors and symbols show which images were processed prior to upload based on either searching (black crosses); or on the basis of the IF; those selected for pre-processing (blue dots), and those which were not (orange crosses). As can be seen and expected there is one peak (the central one) where more images should optimally have been scheduled for pre-processing prior to upload. That they were not is a combination of the heuristics in the sampling strategy, and the uploading speed. That is, they were simply uploaded before the IF (the spline estimate) was good enough to schedule them for pre-processing. The blue line in Figure 7 corresponds to the final spline.

**Figure 7.**
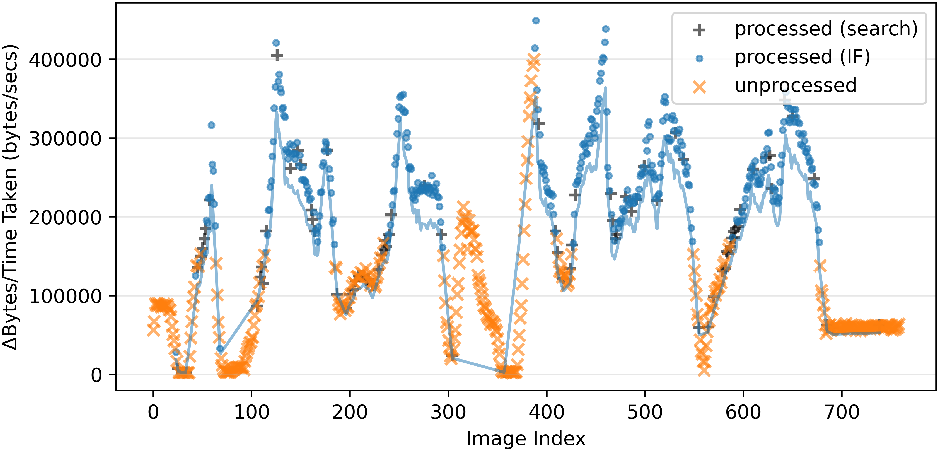
Image size reduction (normalized by CPU cost) over index, showing which images are processed at the edge. Those marked ‘processed’ were processed at the cloud edge prior to upload (and vice-versa) – selected either to *search* for new areas of high/low reduction, or to exploit known areas (using the IF). The line shows the final revision of the splines estimation of the message size reduction (the IF). Note how this deviates from the true value (measured independently for illustration purposes on the same hardware), in regions of low reduction. Note the oscillating pattern which is an artifact movement over the grid in the miniTEM. Adapted from [25].

## Discussion

This paper has discussed an approach to the design and development of smart systems for processing large data streams. The key idea of a HASTE pipeline is based on prioritization with an *interestingness function*, and the application of a *policy*. We demonstrated in two distinct case studies that this simple model can yield significant performance gains for data-intensive experiments. We argue that IFs (and the prioritization and binning that they achieve) should be considered more a ‘first class citizen’ in the next generation of workflow management systems, and that the prioritization of data using IFs and policies are useful concepts for designing and developing such systems.

The ability to express informative IFs are critical to the efficiency of a HASTE pipeline. IFs are chosen by the domain expert to quantify aspects of the data to determine online prioritization. In this work we provide two examples of increasing complexity. In Case Study 1, the IF is a static, idempotent function of a single image – which can be checked against a static threshold, to determine a priority ‘bin’ or tier to store the image. In Case Study 2, the prioritization of the queue of images waiting to be uploaded is revised online, as the underlying model is revised. The strength of our proposed model is that, having defined an IF, by making small changes to the policy, the user is able to reconfigure the pipeline for different deployment scenarios and datasets, with different resulting resource allocation. The HASTE toolkit is an initial implementation of this vision. An avenue for future work will explore the creation of IFs through training in real-time, using active learning and potentially also reinforcement learning.

The policy-driven approach of resource prioritization proposed under the HASTE pipeline paradigm can be generalized to optimize utilization of different forms of constrained computational resources. In some contexts (such as Case Study 1) we are concerned with processing data streams for long-term storage, so storage requirements (and associated costs) are the key concern. In other contexts, with a focus on real time control, automation and robotics, the priority can be more about achieving complex analysis with low-latency. In Case Study 2 this is manifest as a need to achieve edge to cloud upload in the shortest possible time.

The different policies for the two case studies reflect this: in Case Study 1, the user defines a policy to ‘bin’ images according their interestingness score (i.e. image quality), these thresholds are pre-defined by the user. That is to say, the user decides explicit interestingness thresholds, and this determines the resources (in this case, storage) which are allocated, and the final cost. In similar deployment scenarios where cloud storage is used (especially blob storage) costs would depend on the number of images within each interestingness bound. Whereas in Case Study 2, by modelling the predicted extent of message size reduction as an IF within the HASTE tools, we can define a policy to prioritize image processing and upload with the goal of minimizing the total upload time for the next step in the pipeline.

These policies induce two forms of DH: In Case Study 2, the DH is manifest as a priority queue, updated in real time as new images arrive, are pre-processed, and eventually removed - whereas the available resources (CPU, network) are fixed. By contrast, the data hierarchy in Case Study 1 is static, defined by fixed thresholds on interestingness score – in this case, it is the resources (in this case, storage, and consequent processing) which are variable, determined by how many images end up in each tier of the hierarchy.

Finally we note that the IF and policy could also be used to prioritize data based on some measure of confidence. In many scientific analyses there exists a significant amount of uncertainty in several steps of the modeling process. For example in a classification setting the class labels predicted can be highly uncertain. If in the top tier of the hierarchy we would place only those data points for which we are confident in the predicted label, downstream analysis would see a reduction in noise and an increased separability of the (biological) effects under study, as discussed in [27].

## Conclusion

In this paper we have proposed a new model for creating intelligent data pipelines, and presented a software implementation, the HASTE Toolkit. We have shown how these tools can be leveraged in imaging experiments to organize datasets into DHs. We have shown benefits in terms of cost reduction and performance improvement, in terms of compute resources of various kinds). In our case studies, we have studied some typical deployment scenarios, and shown how prioritization can be achieved in these scenarios. Conceptualizing data analysis pipelines around IFs allows better use of various computing resources, and provides a conceptual structure for us to think about the involvement of humans in such pipelines (and their monitoring), as well as a means of managing scientific experimentation - either with instruments or through simulation.

The proposed HASTE pipeline model is intended as a means of bringing structure to large scientific datasets - a means of curatinga *DataLake* [28], whilst avoiding creating a *data swamp* [29, 30]. It is effectively a design pattern creating an data hierarchy from “runtime knowledge” about the dataset – extracted in real time. The HASTE Toolkit is intended to help scientists achieve this.

The key contribution made by the HASTE Toolkit is the design of an API which allows the user to express how they would like their data to be prioritized, whilst hiding from them the complexity of implementing this behaviour for different constrained resources in different deployment contexts. Our hope is that the toolkit will allow intelligent prioritization to be ‘bolted on’ to new and existing systems - and is consequently intended to be usable with a range of technologies in different deployment scenarios.

In the general context of big data, the HASTE Toolkit should be seen an effort to address challenges related to data streams and efficient placement and management of data. It provides the technical foundation for automatically organizing incoming datasets in a way that makes them self-explainable and easy to use based on the features of data objects rather than traditional metadata. It also enables efficient data management and storage based on data hierarchies using dynamic policies. This lays the foundation for domain experts to efficiently select the best-suited data from a massive dataset for downstream analysis.

## Declarations

## List of abbreviations

DH: Data Hierarchy. Conceptual structures in datasets, realized as, e.g. tiered storage systems.
HASTE: Hierarchical Analysis of Spatial (TE)mporal data.
HSC: Haste Storage Client. A core HASTE component for managing data hierarchies.
IF: Interestingness function. Applied to a document in HASTE to compute an interestingness score.
PLLS: Power Log Log Slope.

## Ethical Approval

Not applicable.

## Consent for Publication

Not applicable.

## Acknowledgements

Thanks to Anders Larsson and Oliver Stein for help with software deployment and testing for Case Study 1. Thanks to Polina Georgiev for providing the images used in the evaluation of Case Study 1. Resources from The Swedish National Infrastructure for Computing (SNIC) [31] were used for Case Study 2.

## Footnotes

We use the generic term data object but note that analogous terms in various contexts include: documents, messages and blobs

HASTE: Hierarchical Analysis of Spatial and Temporal Data http://haste.research.it.uu.se/

At the time of writing, supported platforms are: OpenStack Swift, Pachyderm [18], and POSIX-compatible filesystems.

Version 3.1.8 was used for this study.

https://github.com/CellProfiler/Distributed-CellProfiler

